# The Slingshot phosphatase 2 is required for acrosome biogenesis during spermatogenesis in mice

**DOI:** 10.1101/2022.09.15.508144

**Authors:** Ke Xu, Xianwei Su, Kailun Fang, Yue Lv, Gang Lu, Waiyee Chan, Zi-Jiang Chen, Jinlong Ma, Hongbin Liu

**Author notes:** These authors contributed equally to this work (Ke Xu, Xianwei Su, Kailun Fang). To whom correspondence should be addressed (Z-J.C.); (J.M.); (H.L.).

## Abstract

The acrosome is a membranous organelle positioned in the anterior portion of sperm head and is essential for male fertility. Acrosome biogenesis requires the dynamic cytoskeletal shuttling of vesicles towards nascent acrosome which is regulated by a series of accessory proteins. However, much remains unknown about the molecular basis underlying this process. Here, we generated *Ssh2* knock-out (KO) mice and show that Slingshot phosphatase 2 (SSH2), a regulator of actin remodeling, is essential for acrosome biogenesis and male fertility. In *Ssh2* KO males, spermatogenesis was arrested at the early spermatid stage with enhanced germ cell apoptosis and the impaired acrosome biogenesis was characterized by defective transport/fusion of proacrosomal vesicles. Moreover, disorganized F-actin structures accompanied by excessive phosphorylation of COFILIN were observed in testes of *Ssh2* KO mice. Collectively, our data reveal a modulatory role for SSH2 in acrosome biogenesis through COFILIN-mediated actin remodeling and the indispensability of this phosphatase in male fertility in mice.

## Introduction

The most common cause of non-obstructive azoospermia is impaired spermatogenesis (i.e., the production of mature spermatozoa), which is behind about 15% of infertility cases in men (Agarwal et al., 2021). In basic studies of mice, spermatogenesis is routinely subdivided into 12 stages (Meistrich & Hess, 2013; Oakberg, 1956), and the process through which post-meiotic spermatids develop into mature spermatozoa is termed “spermiogenesis” (Bao & Bedford, 2016), marked by a series of cellular remodeling events that includes the biogenesis of the acrosome – an acidic, membranous organelle positioned over the anterior part of the sperm nucleus that functions in the fertilization of the egg (Moreno & Alvarado, 2006). The transformation of round spermatids involves the cytoskeletal system, which shuttles proteins and vesicles to the nascent acrosome via dynamic cytoskeletal remodeling orchestrated by various accessory proteins such as the actin-binding protein PROFILIN-3 (Umer et al., 2021).

According to the classical morphological descriptions of acrosome biogenesis that follows four sequential phases (the Golgi phase, cap phase, acrosome phase, and maturation phase) (Clermont & Leblond, 1955), the nascent acrosome is sequentially assembled from so-called proacrosomal vesicles that originate from the Golgi apparatus through the biosynthetic pathway (Khawar, Gao, & Li, 2019). However, recent experimental findings have implicated some other cellular components (e.g. the endocytic machinery) in vesicular trafficking towards the growing acrosome (Berruti & Paiardi, 2015; Y. C. Li et al., 2006), supporting the existence of extra-Golgi supply of proacrosomal vesicles. It is important to note that the role of filamentous actin (F-actin) as a cytoskeletal transport platform for proacrosomal vesicles has been documented (Kierszenbaum, Rivkin, & Tres, 2003b). In addition, several Golgi-associated proteins are known to function in the transport/fusion of proacrosomal vesicles, including GOPC (Yao et al., 2002), PICK1 (Xiao et al., 2009) and GM130 (Han et al., 2017). More recently, the contribution of autophagic machinery in Golgi-derived proacrosomal vesicle trafficking has been suggested by findings in germ cell-specific *Atg7-KO* (H. Wang et al., 2014) and *Sirt1-KO* (C. Liu et al., 2017) mouse models, offering new insights into the molecular mechanisms through which the acrosome develops.

As an actin-binding protein, COFILIN is widely known for its cutting and depolymerizing functions that promote the subsequent remodeling of actin filaments (Wioland et al., 2017), which is required for many actin-driven possesses such as mitosis (Amano, Kaji, Ohashi, & Mizuno, 2002) and organelle trafficking (Cichon et al., 2012). Intriguingly, COFILIN was also found to stimulate actin nucleation at a high COFILIN/actin concentration ratio, which in turn favors actin filament assembly (Andrianantoandro & Pollard, 2006). It follows that the complex modulation of COFILIN activity, consisting of post-translational modifications, protein binding, and redox reactions (Namme, Bepari, & Takebayashi, 2021), enables the proper rearrangement of F-actin organization in response to various extracellular stimuli. Among these regulatory mechanisms, the phosphorylation/dephosphorylation at the Ser-3 residue stands out as the most crucial. Phosphorylation-mediated inactivation by LIM domain kinases (LIMKs) prevents COFILIN’s actin remodeling functions (Arber et al., 1998), whereas activating dephosphorylation of COFILIN is catalyzed by phosphatases such as chronophin (Gohla, Birkenfeld, & Bokoch, 2005) and the Slingshot (SSH) family of proteins (Niwa, Nagata-Ohashi, Takeichi, Mizuno, & Uemura, 2002). A previous study with respect to mouse reproduction demonstrated that impaired COFILIN phosphorylation results in increased germ cell apoptosis in murine testes (Takahashi, Koshimizu, Miyazaki, & Nakamura, 2002), implicating COFILIN phospho-regulation in the progression of spermatogenesis.

The ssh gene was originally identified in Drosophila mutants with the bifurcation phenotype of bristles and hairs (Niwa et al., 2002), and the mammalian homologs encode a set of COFILIN phosphatases, namely the SSH phosphatases. Some biological relevance of SSHs has been revealed, especially the most well-characterized isoform SSH1 (Bielig et al., 2014), while much less is known about SSH2 other than its role as a regulator of actin remodeling during neutrophil chemotaxis (Xu et al., 2015). According to a gene-expression study focusing on the SSHs, relatively higher expression of SSH2 was observed in testes compared to other organs in mice (Ohta et al., 2003). Moreover, in the light of a recent proteomic study which is aiming at the prediction of meiosis-essential genes, SSH2 exhibited increasing protein abundance changes with the proceeding of murine spermatogenesis (Fang et al., 2021), illustrating its potential role in male germ cell development. By generating mice carrying targeted knockout (KO) of *Ssh2*, we found that SSH2 is an activator of COFILIN-mediated actin cytoskeleton remodeling and is essential for proper acrosome biogenesis during spermatogenesis and thus male fertility. The impaired acrosome biogenesis in *Ssh2* KO mice is likely a consequence of abnormal vesicular trafficking/fusion that can be attributed to disrupted F-actin remodeling.

## Materials and Methods

### Animals

The mouse *Ssh2* gene (Ensembl: ENSMUSG00000037926) is 243.93 kb and is located on chromosome 11, and 15 exons have been identified. The CRISPR/Cas9 genome editing system (Cyagen Biosciences, Suzhou, China) was used to generate *Ssh2* KO mice in the C57BL/6J background. Briefly, single guide RNAs (sgRNAs) targeting the mouse *Ssh2* gene were co-injected with the Cas9 mRNA into fertilized mouse eggs, resulting in the deletion of the genomic DNA segment harboring exon 8 of *Ssh2*. Founder mice were identified by PCR followed by sequence analysis to confirm the deletion, and these mice were crossed with wildtype (WT) mice to test germline transmission and F1 animal generation. All animal-breeding work was carried out at the Laboratory Animal Center of Shandong University where the mice were housed under controlled environmental conditions with free access to pathogen-free water and food. Mice were maintained and collected for experiments according to protocols approved by the Animal Ethics Committee of the School of Medicine, Shandong University. All animal care protocols in this study were reviewed and approved by the Animal Use Committee of the School of Medicine, Shandong University.

### Genotyping

Genomic DNA extracted from mouse tail tips (H. Liu et al., 2019) was subjected to PCR performed in a thermal cycler (T100, Bio-Rad Laboratories, Hercules, CA, USA). The WT and *Ssh2* KO alleles were assayed by primers: 5’-TCC TTG CCT TGA GAA TTC AAG CAA G-3’ (forward) and 5’-TGA TAT GGT TAG TCC ATT GTG CCC A-3’ (reverse). The PCR conditions were as follows: 94°C for 3 min; 35 cycles of 94 °C for 30 sec, 66 °C for 35 sec, and 72 °C for 35 sec; and 72 °C for 5 min. The PCR products were assessed on 2% agarose gels.

### Antibodies and reagents

The rabbit anti-SSH2 polyclonal antibody against amino acids 1,295–1,423 of the mouse SSH2 protein was custom-generated by Dia-an Biological Technology Incorporation (Wuhan, China) as previously described (M. Li et al., 2019). The customized SSH2 antibody was used for western blotting at a dilution of 1:1,000 and for immunofluorescent staining at a dilution of 1:200. The antibodies for western blotting were rabbit anti-COFILIN monoclonal antibody (1:1,000 dilution, 5175, Cell Signaling Technology, Danvers, MA, USA), rabbit anti-phospho-COFILIN monoclonal antibody (1:1,000 dilution, 3313, Cell Signaling Technology), mouse anti-LIMK-1 monoclonal antibody (1:100 dilution, sc-515585, Santa Cruz Biotechnology, Dallas, TX, USA), mouse anti-LIMK-2 monoclonal antibody (1:100 dilution, sc-365414, Santa Cruz Biotechnology), mouse anti-F-actin monoclonal antibody (1:500 dilution, ab130935, Abcam), rabbit anti-BCL2-associated X protein (BAX) antibody (1:1,000 dilution, 2772, Cell Signaling Technology), rabbit anti-BCL2-associated agonist of cell death (BAD) antibody (1:1,000 dilution, 9292, Cell Signaling Technology), rabbit anti-Caspase-3 antibody (1:1,000 dilution, 9662, Cell Signaling Technology), rabbit anti-Cleaved Caspase-3 monoclonal antibody (1:1,000 dilution, 9664, Cell Signaling Technology), rabbit anti-B-cell lymphoma-2 (BCL-2) polyclonal antibody (1:1,000 dilution, 26593-1-AP, Proteintech, Rosemont, IL, USA) and mouse anti-GAPDH monoclonal antibody (1:10,000 dilution, 60004-1-lg, Proteintech). HRP-conjugated Affinipure goat anti-mouse and anti-rabbit IgG (H+L) (SA00001-1/SA00001-2, Proteintech) were used as the secondary antibodies for western blotting. The antibodies for immunofluorescent staining were mouse anti-p-COFILIN monoclonal antibody (1:100 dilution, sc-271921, Santa Cruz Biotechnology), mouse anti-GM130 monoclonal antibody (1:100 dilution, 610822, BD Biosciences, San Jose, CA, USA), rabbit anti-GOPC polyclonal antibody (1:100 dilution, ab37036, Abcam, Cambridge, UK), rabbit anti-LC3 polyclonal antibody (1:200 dilution, 4108, Cell Signaling Technology), rabbit anti-SYCP1 polyclonal antibody (1:500 dilution, ab15090, Abcam), and mouse anti-SYCP3 monoclonal antibody (1:200 dilution, ab97672, Abcam). The secondary antibodies used for immunofluorescent staining were Alexa Fluor 488/594-conjugated goat anti-rabbit IgG (H+L) (A-11008/A-11012, Invitrogen, Carlsbad, CA, USA) and goat anti-mouse IgG H&L (Alexa Fluor 488/594) (ab150117/ab150120, Abcam). Rhodamine phalloidin was used to visualize F-actin (R415, Invitrogen), and Alexa Fluor 488-conjugated lectin PNA from *Arachis hypogaea* was used to visualize acrosomes (L21409, Invitrogen).

### In vivo fertility assessment

For fertility analysis, each adult male (n = 3, 8–10 weeks old) of the different genotypes was mated with two or three age-matched C57BL/6J female mice for 3 months. Females were checked for the presence of vaginal plugs. The number of live-born pups in each cage was counted individually.

### Tissue collection, histological analysis, and immunofluorescence

For histological examination, at least three adult mice for each genotype were analyzed. Testes and caudal epididymides were dissected immediately following euthanasia, fixed in Bouin’s solution (HT1013, Sigma-Aldrich, St. Louis, MO, USA) for 16–24 h at room temperature, dehydrated in an ethanol series (70%, 85%, 90%, 95%, and absolute ethanol), cleared in xylene, and embedded in paraffin. For immunofluorescent staining, testes and epididymides were fixed in 4% paraformaldehyde (PFA, P1110, Solarbio, Beijing, China) overnight at 4°C, dehydrated, cleared, and embedded. The tissues were then cut into 5 μm sections using a microtome (HistoCore BIOCUT, Leica Biosystems, Nussloch, Germany) and coated onto glass slides. Following drying at 60°C for 1 h, the sections were deparaffinized in xylene, hydrated by a graded alcohol series, and stained with hematoxylin for histological analysis or stained with periodic acid Schiff (PAS)- hematoxylin (ab150680, Abcam) for determining the seminiferous epithelia cycle stages. For TUNEL staining, we followed the manufacturer’s instructions (KeyGen Biotech, Nanjing, China). Images were collected under a microscope with a coupled camera device (BX53, Olympus, Tokyo, Japan). For immunostaining, deparaffinized sections were hydrated, immersed in sodium citrate buffer (pH 6.0) and heated for 15 min in a boiling water bath for antigen retrieval. After permeabilization with PBS containing 0.3% Triton X-100 and blocking with 5% bovine serum albumin, the slides were incubated with primary antibodies overnight at 4°C. Then the sections were rinsed in PBS and incubated with appropriate FITC-conjugated secondary antibodies and/or Alexa Fluor 488-conjugated PNA/rhodamine phalloidin for 60 min at room temperature. Mounting medium with DAPI (ab104139, Abcam) was used to visualize the nucleus and to mount the slides.

Immunostaining was also conducted on cryosections that were prepared with the following procedure. Fresh tissues were fixed in 4% PFA for 12–24 h at 4°C, dehydrated in 20% sucrose (in 1× PBS) for 2–3 h followed by 30% sucrose (in 1× PBS) overnight at 4°C, embedded in OCT, and cut into cryosections at 8 μm thickness using a cryotome (CM1950, Leica Biosystems). Once sectioned, the samples were fixed on the slides with 4% PFA for 10–15 min at room temperature and washed three times with PBS. Subsequent steps were performed in the same way as for staining paraffin sections.

### Sperm count

Mice epididymal sperm were released into PBS through multiple incisions of the cauda followed by incubation for 30–60 min at 37°C under 5% CO_2_. The living sperm was then diluted in PBS at 1:50 and transferred to a hemocytometer for counting. For all experiments, adult males 2–6 months of age were used.

### Protein extraction and western blot analysis

Testes from adult mice were cut into ~20 mg pieces and transferred into ~200 μl pre-cooled denaturing buffer from the Minute Total Protein Extraction Kit (SD-001, Invent Biotech, Plymouth, MN, USA) containing protease inhibitors (CO-RO, Roche, Indianapolis, IN, USA) and phosphatase inhibitors (PHOSS-RO, Roche). Protein extraction was performed with the Minute Total Protein Extraction Kit following the manufacturer’s instructions. In order to load equal amounts of protein, the protein concentration in the supernatant was measured using the Pierce BCA Protein Assay Kit (23225, Thermo Fisher Scientific, Waltham, MA, USA). After diluting with loading buffer and boiling for 10 min at 95°C, an aliquot of 20 mg protein lysate per sample was resolved by sodium dodecyl sulfate-polyacrylamide gel electrophoresis, transferred to polyvinylidene fluoride membranes, blocked with noise-cancelling reagents (WBAVDCH01, Millipore, Burlington, MA, USA), and immunoblotted with the appropriate primary antibodies. The membranes were then incubated with HRP-conjugated secondary antibodies followed by immunodetection using a chemiluminescence imaging system (5200, Tanon Technology, Shanghai, China).

### Surface chromosome spreading

Chromosome spread analysis was conducted with the drying-down technique as previously described (Peters, Plug, van Vugt, & de Boer, 1997). Briefly, spermatocytes from hypotonicity-treated testicular tubules were suspended in 0.1 M sucrose, spread on glass slides, and immersed in PFA solution containing 0.15% Triton X-100. Subsequent immunolabeling of spermatocyte nuclei using antibodies against SYCP1 and SYCP3 was performed according to the protocol for immunofluorescent staining of cryosections.

### Isolation of spermatogenic cells and fluorescence-activated cell sorting (FACS)

Spermatogenic cells were isolated from the testes of 8-week-old mice and subsequently subjected to flow cytometry and fluorescence-activated cell sorting (FACS) for DNA ploidy analysis. In brief, testes collected from 8-week-old mice were decapsulated and washed in pre-chilled PBS three times. After carefully cutting, pieces of the seminiferous tubules were transferred into PBS containing 120 U/ml collagenase I (17018029, Gibco, Carlsbad, CA, USA) for a 5 min incubation at 32°C. To obtain cell suspensions, the tubular pieces were incubated in 0.25% trypsin (25200056, Gibco) containing 1 mg/ml DNase I (18047019, Thermo Fisher Scientific) at 32°C for 8 min with gentle pipetting.

Cold DMEM containing 5% fetal bovine serum (FBS, 10091155, Gibco) was added to stop the digestion. The spermatogenic cells were collected from the cell suspension after filtration through PBS-saturated 70 μm cell filters followed by centrifugation at 4°C (500 × *g* for 5 min). After removal of the supernatant, the cells were resuspended in 1 ml DMEM containing 2 μl Hoechst 33342 (62249, Thermo Fisher Scientific), 2 μl Zombie Aqua dye (423101, BioLegend, San Diego, CA, USA), and 5 μl DNase I and subjected to flow cytometry. For FACS, the cell suspension was further rotated at a speed of 10 rpm/min for 20 min at 32°C and centrifuged for 5 min at 4°C. Finally, the cells prepared for sorting were resuspended and cell populations were collected based on their fluorescent label with Hoechst 33342 staining using FACS.

### Electron microscopy

For transmission electron microscopy, testes collected from adult mice were cut into 1–2 mm^3^ pieces and fixed in 2.5% glutaraldehyde in 0.1 M cacodylate buffer (pH 7.4) at 4°C overnight. After washing in 0.1 M cacodylate buffer three times, the samples were incubated in 1% osmium tetroxide at room temperature for 1 h. The samples were then washed with ultrafiltered water and stained in 2% uranyl acetate for 30 min. Subsequent dehydration was done through consecutive incubation in graded ethanol series (50%, 70%, 90%, and absolute ethanol) and an acetone bath. After dehydration, samples were sequentially incubated in Embed-812 resin:acetone mixtures at 1:1 and 3:1 for 2 h each and in 100% Embed-812 resin for 3–4 h, all at room temperature. The samples were then orientated and embedded with fresh resin at 37°C. Tissue blocks were cut on an ultramicrotome (UC7, Leica Biosystems) to yield 80 nm ultrathin sections, and these were subsequently contrasted with uranyl acetate and lead citrate. The sections were then observed and imaged using a transmission electron microscope operating at 120 kV (JEM-1400, JEOL, Pleasanton, CA, USA).

### Mouse seminiferous tubule squashes

Seminiferous tubule squash slides were prepared as described previously (Wellard, Hopkins, & Jordan, 2018) with minor modifications. Specimens of testes were removed from adult mice and washed with PBS. The testicular tunica albuginea was torn and the seminiferous tubules were released, collected, and fixed with 2 ml fixing solution for 5 min at room temperature in a 35 mm Petri dish. After washing twice with PBS, the seminiferous tubules were cut into small pieces (about 10–20 mm in length) that were subsequently transferred to the prepared glass slides containing 100 μl fixing/lysis solution. Coverslips were then applied to cover the glass slides and were compressed tightly for 20 seconds. After squashing, the glass slides were immediately frozen in liquid N_2_ for 15 sec and stored at −80°C. To perform immunolabeling on the squash slides, the coverslips were first removed after rewarming. The slides were immersed in PBS and washed three times. The subsequent steps were similar to the procedure for immunofluorescent staining of cryosections.

### Imaging

Immunolabeled slides, including paraffin-sections, cryosections, squash slides, and chromosome spreads, were imaged by confocal microscopy (Dragonfly Spinning Disc confocal microscope driven by Fusion Software, Andor Technology, Belfast, UK). Projection images were then processed and analyzed using Bitplane Imaris (version 9.7) software. The histological and terminal deoxynucleotidyl transferase-dUTP nick-end labeling (TUNEL)-stained samples were imaged using an epifluorescence microscope (BX52, Olympus) equipped with a digital camera (DP80, Olympus) and processed using cellSens Standard (Olympus) software packages.

### Statistical analysis

Statistical analyses were conducted with SPSS software (version 22.0, IBM Corporation, Armonk, NY, USA). All data are presented as the mean ± SEM, as indicated in the figure legends. The statistical significance of the difference between the mean values for the various genotypes was determined by Welch’s t-test with a paired two-tailed distribution. The data were considered significant when the P-value was less than 0.05.

## Results

### *Ssh2* is essential for male fertility

To explore the *in vivo* functions of SSH2, we generated mice with *Ssh2* KO lacking exon 8 using CRISPR/Cas9 genome editing (Figure 1A). We validated the *Ssh2* KO in mice using PCR of tail-derived genomic DNA (supplementary Figure S1) and western blotting of whole testicular lysates. SSH2 was immunodetected in the testes of wild-type (WT) mice but was absent in the *Ssh2* KO samples (Figure 1B). *Ssh2* KO male mice were found to be completely infertile (Figure 1C). Six mice of each genotype were assessed for breeding over a 2-month period, while no pups were obtained when adult *Ssh2* KO males mated with 8-week-old WT fertile females. We detected no differences in the size of testis, body weight, testis weight, or testis-to-body weight ratio in *Ssh2* KO male mice compared to their WT littermates (Figure 1D-G).

**Fig. 1.**
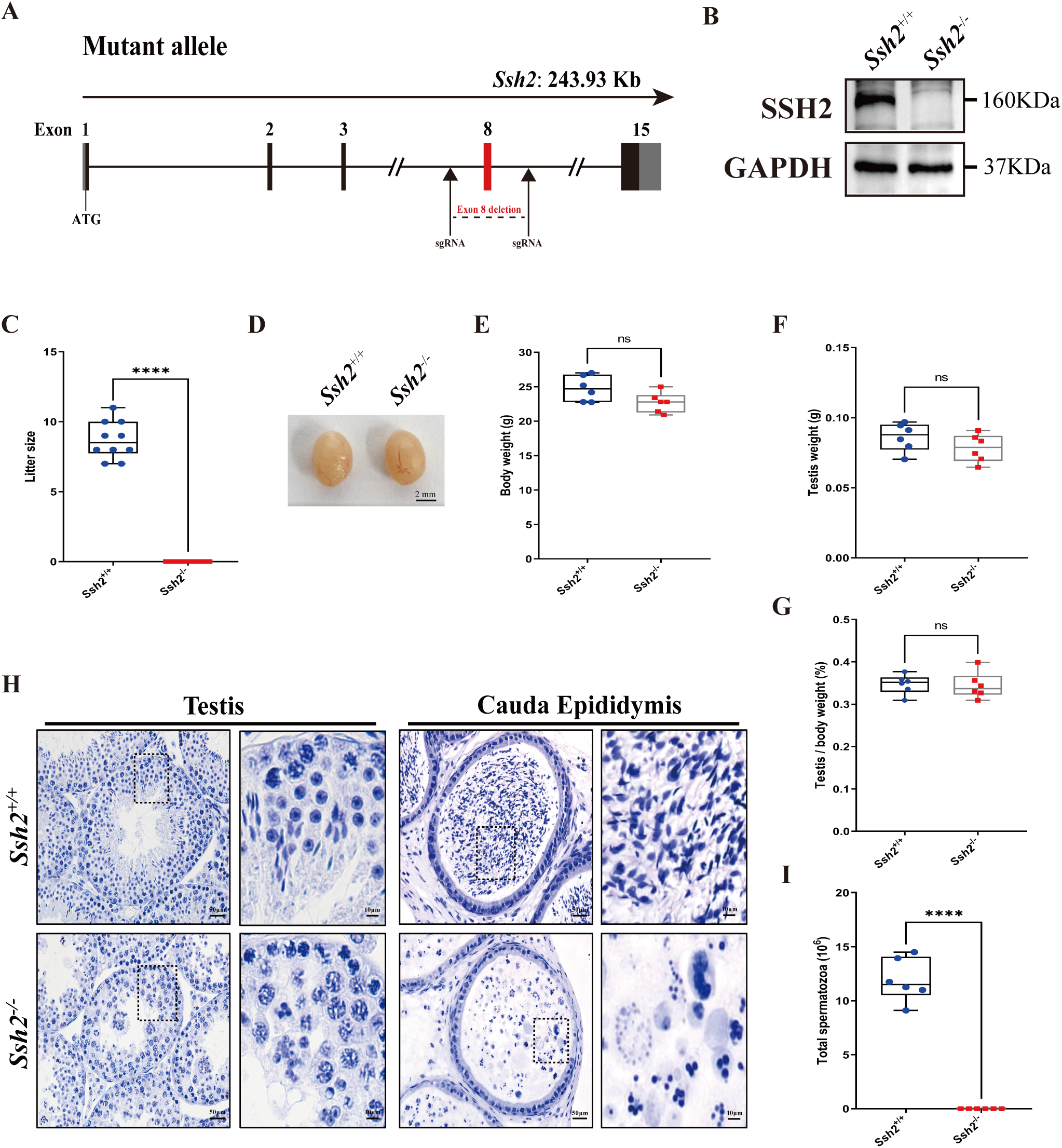
*Ssh2* KO causes severe reproductive defects and male infertility in mice. **(A)** Schematic representation of the generation of *Ssh2* KO mice using CRISPR/Cas9. **(B)** Validation of *Ssh2* KO by western blotting in whole testis lysates from WT and *Ssh2* KO 8-week-old mice (n = 3), indicating the absence of SSH2 protein in *Ssh2* KO testes. GAPDH was used as the loading control. **(C)** Number of pups per litter from WT (8.70 ± 0.42) and *Ssh2* KO (0.00) male mice (8 weeks old) after crossing with WT female mice (8–10 weeks old) for 3 months (n = 10). Data are presented as the mean ± SEM; ****p < 0.0001, calculated by Student’s t-test. Bars indicate the range of data. **(D)** The testes from *Ssh2* KO mice appeared phenotypically normal when compared to testes of WT mice at 8 weeks of age, n = 6. Scale bars: 2 mm. **(E-G)** Body weights, weights of the testes, and the testis-to-body weight ratio of WT and *Ssh2* KO males at 6–8 weeks of age (n = 6). Data are presented as the mean ± SEM; p > 0.05 calculated by Student’s t-test. Bars indicate the range of data. **(H)** Histology of the testis and cauda epididymis from WT and *Ssh2* KO mice. Sections were stained with hematoxylin. No elongating or elongated spermatids were detected in *Ssh2* KO testes, and no mature sperm were detected in the *Ssh2* KO epididymis. Boxed regions are magnified on the right. Scale bars: 50 μm for the original region (left columns); 10 μm for the magnified region. Images are representative of testes/cauda epididymis extracted from at least six adult male mice per genotype. **(I)** Total sperm number in the cauda epididymis of WT (11.92 × 106 ± 0.82 × 106) and *Ssh2* KO (0.00) mice, n = 6. Data are presented as the mean ± SEM; ****p < 0.0001, calculated by Student’s t-test. Bars indicate the range of data. **Source data 1.** Observational datasets and original blots.

Histological examination of WT and *Ssh2* KO testes and epididymides by hematoxylin staining revealed impaired spermatogenesis in *Ssh2* KO males, with no mature spermatozoa observed in the epididymal lumen of *Ssh2* KO mice (Figure 1H), which was confirmed by the sperm count in the cauda epididymis (Figure 1I). The seminiferous tubules of WT testes were full of spermatogenic cells and contained a basal population of spermatogonia, spermatocytes, and spermatids. In contrast, almost no elongating or elongated spermatids were observed in the testes of *Ssh2* KO males; instead, round spermatids were found to aggregate to form giant multinucleated cells that were then verified to be spermatid clusters by FACS assessment (Figure 1H, supplementary Figure S2). Thus, we deduced from these findings that *Ssh2* is essential for male fertility.

### *Ssh2* KO mice exhibit spermatogenic arrest at the early spermatid stage and display aberrantly high germ-cell apoptosis

To determine the time point of onset of spermatogenic arrest in *Ssh2* KO males, we first analyzed spermatocyte development by examining chromosomal synapsis during meiotic progression in chromosome spreads from 3-week-old testes of WT and *Ssh2* KO mice. By immunofluorescence co-staining of synaptonemal complex protein 1 (SYCP1) and SYCP3 in the nuclei of spermatocyte spreads – which comprise the central and lateral elements of the synaptonemal complex, respectively (de Vries et al., 2005; Yuan et al., 2000) – we found no obvious differences in their distribution patterns in various stages of meiotic prophase, indicating that the prophase I process was successfully completed in WT and *Ssh2* KO mice (supplementary Figure S3).

We then performed detailed histological analysis by hematoxylin staining of testicular sections from WT and *Ssh2* KO males at various developmental stages. Testes collected from *Ssh2* KO mice at postnatal day (PD)7 and PD14 were indistinguishable from those of WT mice. The most advanced cells in WT testes were elongating spermatids at PD28, and none of these were present in *Ssh2* KO testes. Round spermatid clusters were first observed in *Ssh2* KO tubules at PD28, and these accumulated from PD35 to PD60. We next examined PD35 testes, and histological images showed that the *Ssh2* KO tubules were devoid of elongated spermatids (supplementary Figure S4).

To determine at which stage of spermatogenesis the developmental defects occurred, sections of WT and *Ssh2* KO testes were stained with hematoxylin and PAS, which labels acrosomal glycoproteins, and we observed that spermatogenesis of *Ssh2* KO mice was arrested at stage II-III (Figure 2A). Spermatids at various steps of spermiogenesis were observed in WT seminiferous tubules, whereas only step 1–3 round spermatids were present in *Ssh2* KO tubules, indicating spermiogenic failure in these mice. We also detected round spermatids with malformed acrosomal structures in *Ssh2* KO testes, which suggests that the acrosome formation was also affected. Furthermore, clustered round spermatids and apoptotic-like germ cells exhibiting small hyperchromatic nuclei and polynuclear structures, which are known indicators of apoptosis (Catalano-Iniesta et al., 2019), were observed in *Ssh2* KO tubules (Figure 2A).

**Fig. 2.**
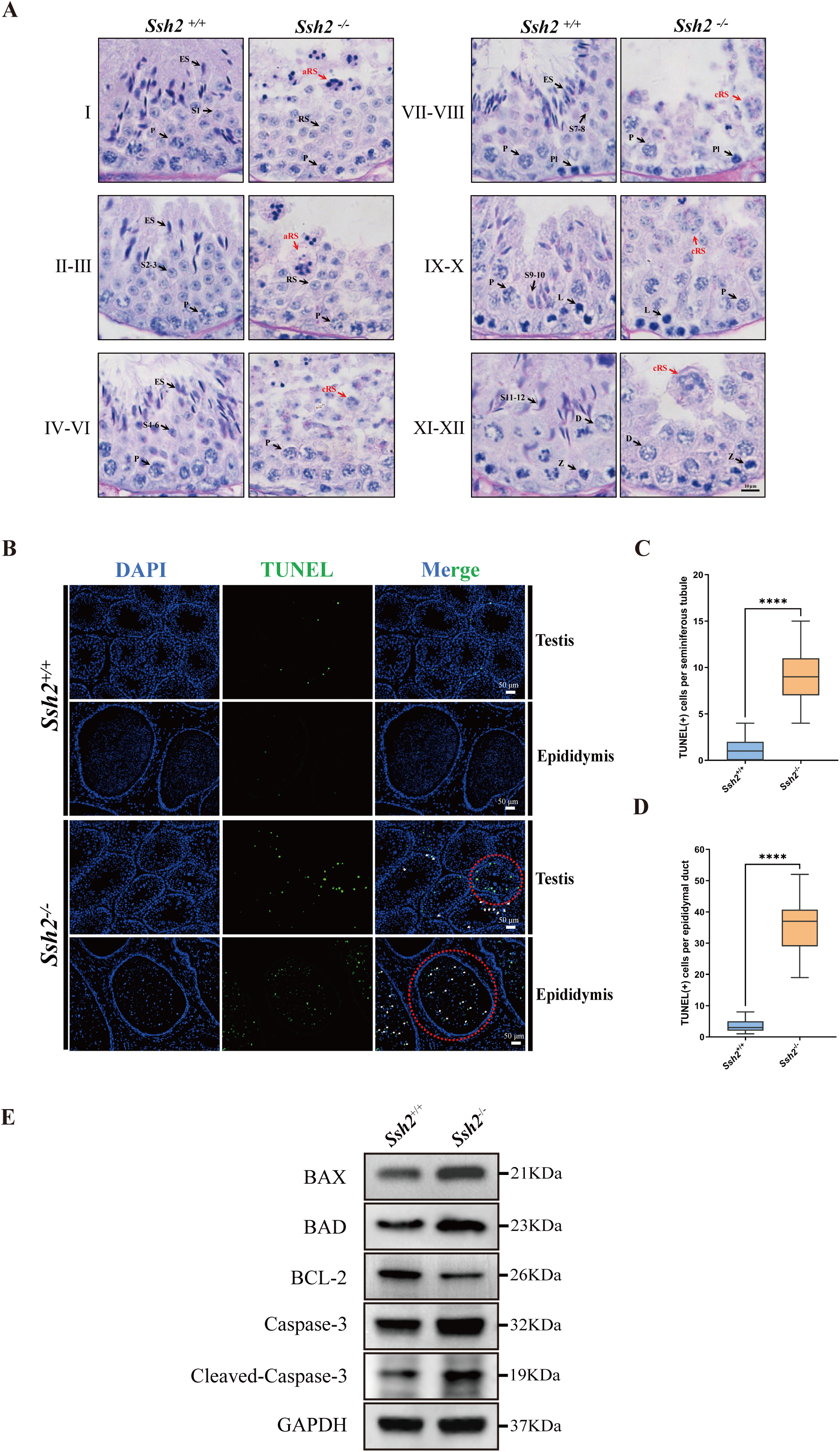
Spermatogenic arrest and enhanced germ-cell apoptosis result from *Ssh2* KO. **(A)** PAS-hematoxylin–stained sections of seminiferous epithelia from WT mice and *Ssh2* KO mice (8 weeks old, n = 3), indicating spermatogenic arrest at steps 2–3 of spermatogenesis in the *Ssh2* KO mice. L, leptotene spermatocytes; D, diplotene spermatocytes; Z, Zygotene spermatocytes; P, pachytene spermatocytes; RS, round spermatids; ES, elongated spermatids; S1–S12, spermatids at different spermiogenic steps; aRS: apoptotic-like round spermatids; cRS: clustered round spermatids. Scale bar: 10 μm. **(B)** TUNEL immunofluorescence staining of the testicular and epididymal sections from WT and *Ssh2* KO mice. The TUNEL-positive puncta and signal intensity significantly increased in *Ssh2* KO testes and epididymides (as the red circles indicate). Green: TUNEL-positive signal; Blue: DAPI; White arrows: TUNEL-positive germ cells. Scale bars: 50 μm. At least three mice (6–8 weeks old) of each genotype were used in the analysis. **(C)** Counts of TUNEL-positive cells per seminiferous tubule in adult *Ssh2* KO testes (8.92 ± 0.21) compared with control (1.19 ± 0.08). Three mice of each genotype were assessed and 50 tubules were validated for each mouse. Data are shown as mean ± SEM; ****p < 0.0001, calculated by Student’s t-test. Bars indicate the range of data. **(D)** Counts of TUNEL-positive cells per epididymal duct in adult *Ssh2* KO testes (35.85 ± 1.06) compared with control (3.57 ± 0.21). Three mice of each genotype were assessed and 20 ducts were validated for each mouse. Data are shown as mean ± SEM; ****p < 0.0001, calculated by Student’s t-test. Bars indicate the range of data. **(E)** Immunoblotting analysis of BAX, BAD, BCL-2, Caspase-3 and cleaved Caspase-3 in testicular lysates from adult WT and *Ssh2* KO mice, n = 3; GAPDH was used as the loading control. **Source data 2.** TUNEL-related observational datasets and original images/blots.

We then conducted a TUNEL assay to identify apoptotic cells in the testes and epididymides of WT and *Ssh2* KO mice. The counts of TUNEL-positive cells were significantly increased in the *Ssh2* KO seminiferous tubules which were observed primarily as apoptotic spermatids, as indicated by the nuclear morphology and location within spermatogenic epithelium (Figure 2B, Figure 2C). Similarly, we found large numbers of isolated apoptotic spermatids in the lumen of epididymal ducts of *Ssh2* KO mice (Figure 2B, Figure 2D). To understand the potential mechanism behind the enhanced germ cell apoptosis elicited by *Ssh2* KO, we estimated the expression levels of several apoptotic signaling proteins including Caspase-3 with BCL-2, BAX and BAD, which are well-known as Bcl-2 family members involved in Caspase-3 activation (Gu et al., 2021). As demonstrated by western blotting, in comparison with that of WT mice, remarkably elevated protein levels of pro-apoptotic BAX, BAD, Caspase-3 and cleaved Caspase-3 accompanied by relative decreased protein levels of anti-apoptotic BCL-2 proteins were detected in the testicular lysates of *Ssh2* KO mice (Figure 2E), suggesting the induction of Caspase-3 activation in the spermatogenic cell apoptosis resulted from *Ssh2* KO. Consistently, these findings support the hypothesis that *Ssh2* KO results in round spermatid arrest with malformed acrosomes during spermatogenesis and that this in turn causes spermatid apoptosis via the Bcl-2/Caspase-3 pathway.

### *Ssh2* KO leads to disrupted acrosome biogenesis

To better understand the defects in *Ssh2* KO spermatid development, we performed an immunofluorescence analysis to evaluate the morphology of the differentiating spermatids in WT and *Ssh2* KO mice. Spermatids in distinct phases of acrosome biogenesis were stained with fluorescein-conjugated peanut agglutinin (PNA), a protein that specifically binds to the outer acrosomal membrane (Cheng et al., 1996). We found spermatids of all spermiogenic steps in the testes of WT males: as spermiogenesis progressed, singular acrosomal vesicles derived from proacrosomic vesicles were formed in the Golgi phase (steps 1—3), became flattened during the cap phase (steps 4—6), spread to the laminae (steps 7—8), and were coated onto the nuclei in the acrosome/maturation phase (steps 9— 16) (Figure 3A, upper panel). On the contrary, instead of intact acrosomal vesicles as seen in WT spermatids, multiple scattered proacrosomic vesicles separated from nuclei (without fusing with each other) were present in a majority of the *Ssh2* KO spermatids (step 1—3) (Figure 3A, lower panel), demonstrating that acrosome biogenesis was disrupted starting from the Golgi phase in *Ssh2* KO mice.

**Fig. 3.**
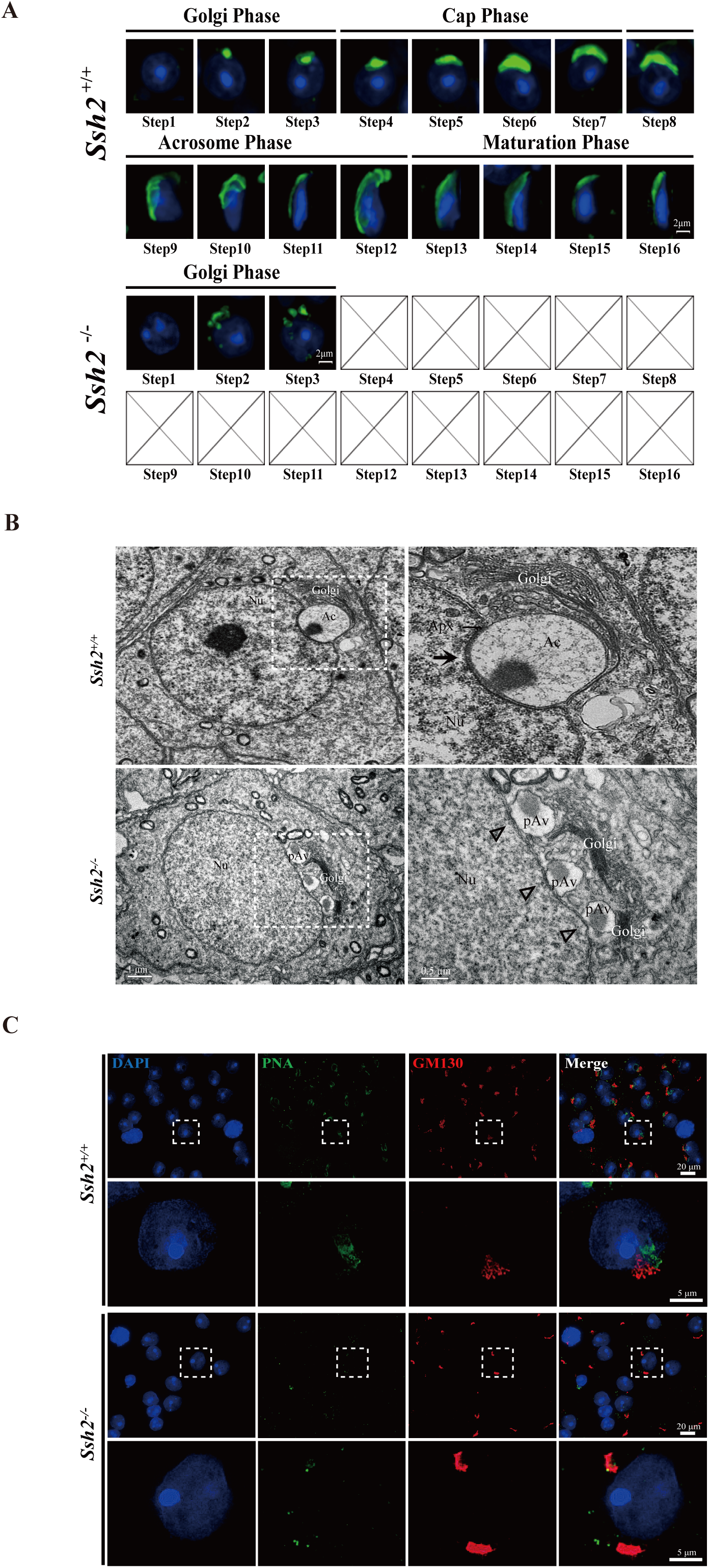
Acrosome biogenesis is disrupted during spermiogenesis in *Ssh2* KO mice. **(A)** Analysis of spermiogenesis and acrosome biogenesis in WT and *Ssh2* KO mice (8 weeks old, n = 3) by fluorescence imaging of spermatids labeled with PNA lectin (green). Nuclei were stained with DAPI (blue). The phases of acrosome biogenesis and corresponding spermiogenesis steps are the Golgi phase (steps 1–3), cap phase (steps 4–7), acrosome phase (steps 8–12), and maturation phase (steps 13–16). Scale bars: 2 μm **(B)** Ultrastructural analysis of WT and *Ssh2* KO spermatids. Intact acrosomes (black arrow) were observed in WT mice. The hollow triangles indicate proacrosomal vesicles that failed to fuse in *Ssh2* KO mice. The regions outlined by white boxes are shown at higher magnification to the right. Nu, nucleus; Ac, acrosome; Golgi, Golgi apparatus; Apx, acroplaxome; Pav, proacrosomal vesicle. Scale bars: left panel, 1 μm; right panel, 0.5 μm. **(C)** Immunofluorescence analysis of the vesicular fusion-related Golgi-specific protein GM130 (red) in WT and *Ssh2* KO round spermatids in testicular sections co-stained with PNA lectin (green). Nuclei were stained with DAPI (blue). Scale bars: original images, 20 μm; magnified images, 5 μm. **Source data 3.** Original images.

### Proacrosomal vesicles do not fuse normally in *Ssh2* KO spermatids

In order to gain insights into the defective acrosome formation in spermatids of *Ssh2* KO mice in the Golgi phase, which is characterized by the fusion of Golgi-derived proacrosomal vesicles (Berruti & Paiardi, 2015; Khawar et al., 2019), we next performed a transmission electron microscopy (TEM) analysis to explore the ultrastructure of the acrosome in spermatids in WT and *Ssh2* KO testes. TEM showed that WT spermatids had a single, large acrosomal vesicle in which the granule was attached to the concave surface of the nucleus and the inner acrosomal membrane was docked to the electron-dense acroplaxome. However, in *Ssh2* KO spermatids, mutiple Golgi-derived small proacrosomal vesicles were adjacent to the trans-face of Golgi apparatuses which contain thick Golgi stacks and failed to fuse together to yield the acrosomal vesicle. Moreover, no acroplaxome-like structure was observed near the nuclear envelope (Figure 3B). These results suggest that the impairment of acrosome formation in *Ssh2* KO mice might result from the failure of proacrosomal vesicle fusion and/or from defects in vesicular trafficking towards the nuclear surface.

Note that these impaired acrosome biogenesis phenotypes appear similar to observations in mice with mutations in *Gm130* (Han et al., 2017), which is known to function in Golgi membrane dynamics and and fusion of Golgi-derived vesicles (Koreishi et al., 2013; Walker et al., 2004). To determine whether *Ssh2* KO affects the fusion of proacrosomal vesicles in the Golgi phase, we performed co-immunostaining of GM130 with PNA staining of seminiferous tubule squashes of WT and *Ssh2* KO mice. We observed GM130 signals with vesicle-like and stack-like shapes in close contact with the PNA-labeled acrosomal vesicles in WT spermatids. However, in *Ssh2* KO spermatids, only large GM130-positive fluorescent aggregates that exhibited no association with PNA-positive proacrosomal vesicles were observed (Figure 3C), suggesting that the localization of vesicular fusion-associated GM130 is altered in Golgi-phase spermatids by *Ssh2* KO. Collectively, these data indicate that *Ssh2* KO causes failure of proacrosomal vesicle fusion.

### SSH2 accumulates at the acrosomal region in round spermatids

To investigate the functions of SSH2 in spermatogenesis, we examined the pattern of SSH2 accumulation in testes at several developmental time points using western blotting. We observed a gradual increase in accumulation during murine spermatogenesis. The protein was undetectable in PD7 testes, became detectable at PD14, and was substantially increased by PD21, the time when round spermatids first appear in seminiferous tubules (Guan et al., 2020), suggesting a potential function of SSH2 in spermatid development (Figure 4A).

**Fig. 4.**
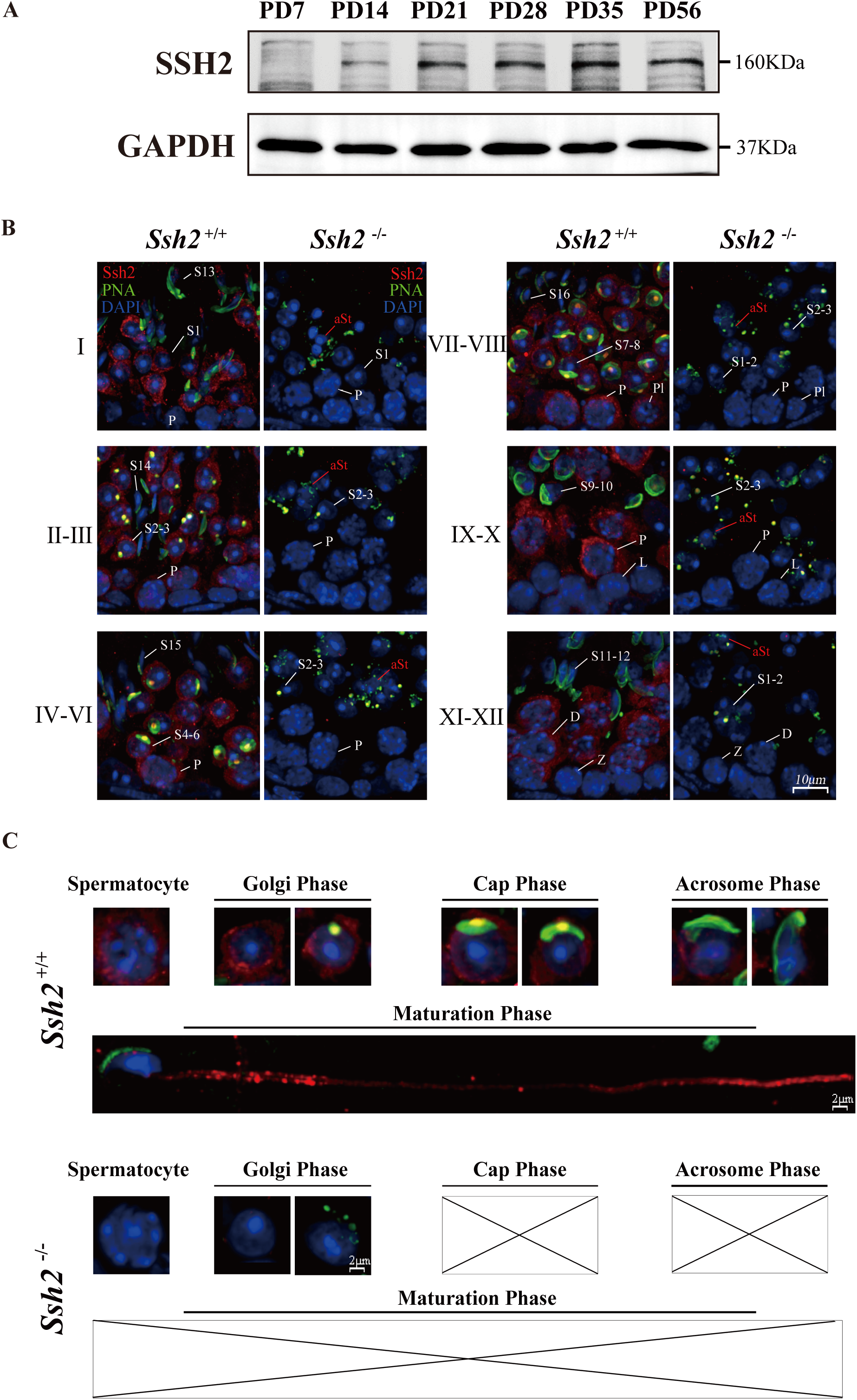
SSH2 accumulates at the acrosomal region of spermatids. **(A)** Immunoblotting against SSH2 in wild-type murine testes sampled at the indicated postnatal days. The expression of Ssh2 at PD14, PD21, PD28, PD35, and PD56 was measured. GAPDH was used as the loading control. **(B)** Co-immunofluorescence staining of Alexa Fluor 488-conjugated PNA lectin (green) and SSH2 (red) on testicular sections from WT (left lane) and *Ssh2* KO (right lane) mice. Nuclei were stained with DAPI (blue). The epithelial spermatogenic cycle is routinely divided into 12 stages on the basis of changes in the morphology of the acrosome and nucleus in spermatids, and this was determined using the combination of PNA lectin and DAPI staining according to the established criteria. Cytoplasmic SSH2 localization in spermatocytes and spermatids of WT mice was observed, while fractured acrosomes were observed at all stages of spermatogenesis in Ssh2 KO mice. P, pachytene spermatocytes; Pl, preleptotene spermatocytes; L, leptotene spermatocytes; Z, zygotene spermatocytes; D, diplotene spermatocytes; S1–16, step 1–16 spermatids. aRS: apoptotic-like spermatids. Scale bar: 10 μm. **(C)** Analysis of acrosomal morphogenesis in spermatogenic cells by co-staining of PNA lectin (green) and SSH2 (red) in WT (upper panel) and *Ssh2* KO (lower panel) murine testes. Nuclei were stained with DAPI (blue). Acrosomal morphology during acrosome biogenesis (Golgi, cap, acrosome, and maturation phase) is shown. No cap, acrosome, or maturation-phase spermatids were observed in *Ssh2* KO mice. Scale bar: 2 μm. **Source data 4.** Original images/blots.

To determine the developmental roles of SSH2 in spermatogenic cells, we performed counterstaining of SSH2 with PNA on seminiferous tubule paraffin sections of WT and *Ssh2* KO mice. SSH2 accumulated predominantly in spermatocytes and post-meiotic round spermatids. In addition to the ubiquitous distribution observed in the cytoplasm of the germ cells, we noted bright, isolated SSH2-positive puncta positioned in the acrosomal region next to the nuclei of round spermatids (Figure 4B, left column). In contrast, fragmented acrosomes with a lack of SSH2 signal were detected in *Ssh2* KO spermatids (Figure 4B, right column).

To further understand the involvement of SSH2 in acrosome biogenesis, we assessed its subcellular localization in spermatids at the different phases. Spermatids in seminiferous tubule squashes obtained from adult WT and *Ssh2* KO mice were immunostained with PNA and SSH2 for further observation in greater detail. Well-formed acrosomes in developing spermatids of WT mice exhibited diverse shapes at the different phases of acrosome biogenesis (Figure 4C, upper panels). In contrast, many irregular, fractured PNA fluorescence signals were observed in most of the *Ssh2* KO spermatids, and no cap-like PNA-positive structures were detected (Figure 4C, lower panels). Moreover, the association of SSH2 with the growing acrosome was clearly evident by the acrosomal localization of SSH2 in the spermatids of WT mice (Figure 4C, upper panels). Together these results show that SSH2 is functionally involved in acrosome biogenesis.

### *Ssh2* KO alters F-actin organization in developing spermatids

It has been reported that SSH2 binds to F-actin in cultured HeLa cells, where it antagonizes LIMK1-induced actin polymerization (Ohta et al., 2003). To test whether F-actin organization in spermatids is affected by *Ssh2* KO during acrosome biogenesis, we first visualized F-actin structures in WT and *Ssh2* KO testicular sections at PD21, PD35, and PD60 using fluorochrome-conjugated phalloidin. As the representative images showed, the F-actin staining was altered in *Ssh2* KO spermatids. Unlike the expected sharp, sickle-shape heads of elongated spermatids surrounded by organized F-actin bundles seen in WT mice, the F-actin aggregated to form numerous lumps in *Ssh2* KO round spermatids (supplementary Figure S5). This aberrant accumulation of F-actin suggests that loss of *Ssh2* accelerates filament growth in spermatids rather than inducing filament disassembly.

To further compare the successive changes in F-actin morphology during acrosome biogenesis in WT and *Ssh2* KO developing spermatids, we performed immunofluorescence microscopy on testicular sections co-stained with phalloidin and PNA. Phalloidin staining was diffusely distributed in the cytoplasm of round spermatids and intact F-actin bundles located around the laminal acrosomes in elongated spermatids of WT mice. In contrast, punctate phalloidin signals accompanied by thick F-actin bundles were mostly dissociated from the fractured acrosomes in *Ssh2* KO spermatids (Figure 5A). Moreover, we co-immunostained the spermatids with phalloidin and PNA in seminiferous tubule squashes from WT and *Ssh2* KO mice. As expected, confocal microscopy showed that punctate F-actin signals were distributed ubiquitously throughout the cytoplasm of Golgi-phase spermatids of WT mice, with relatively intense fluorescence signals localized at acrosomal regions. However, robust phalloidin staining was scattered irregularly in *Ssh2* KO spermatids and exhibited no association with the acrosomal signals (Figure 5B). Together, these findings indicate that F-actin is disorganized in *Ssh2* KO spermatids during acrosome biogenesis.

**Fig. 5.**
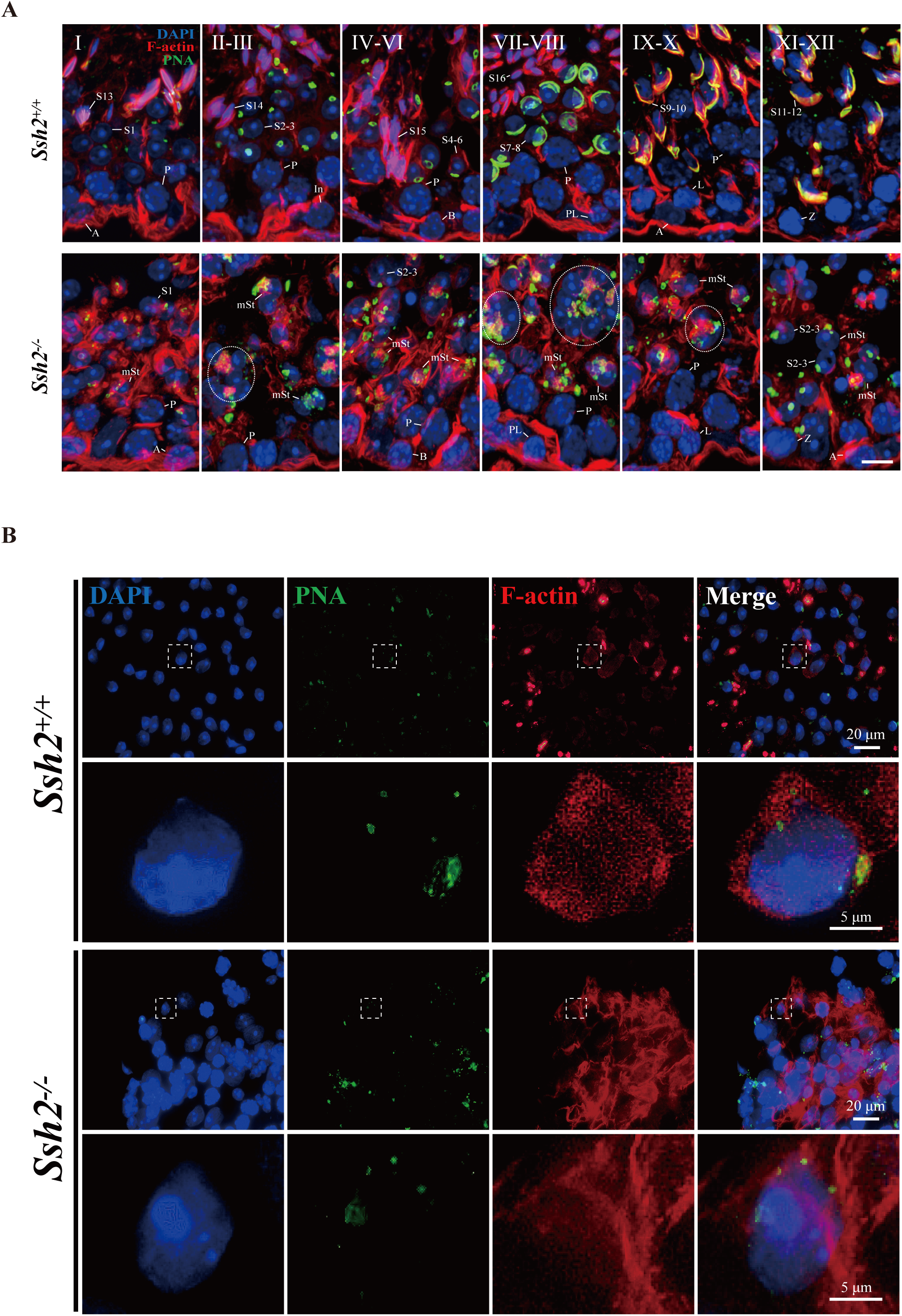
*Ssh2* KO results in disturbed F-actin remodeling in spermatids. **(A)** Immunofluorescence detection of F-actin (red) and PNA lectin (green) during spermatogenesis in testicular sections from WT (top lane) and *Ssh2* KO (bottom lane) mice at PD85. Blocky actin filaments (as indicated by the circles) were observed in spermatids with malformed acrosomes. A, type A spermatogonia; In, intermediate spermatogonia; B, type B spermatogonia; P, pachytene spermatocytes; Pl, preleptotene spermatocytes; L, leptotene spermatocytes; Z, zygotene spermatocytes; D, diplotene spermatocytes; S1–16, step 1–16 spermatids; mSt, spermatids with malformed acrosomes. Scale bar: 10 μm. **(B)** Fluorescence analysis of F-actin (red) and PNA lectin (green) on squashes from WT and *Ssh2* KO mice. Nuclei were stained with DAPI (blue). Diminutive actin filaments exhibited a uniform cytoplasmic distribution in WT spermatids, whereas F-actin was remodeled in *Ssh2* KO spermatids. The framed regions are magnified beneath. Scale bars: original images, 20 μm; magnified images, 5 μm. **Source data 5.** Original images.

### *Ssh2* KO impairs proacrosomal vesicle trafficking

Given the disrupted acrosome biogenesis and disorganized F-actin in *Ssh2* KO spermatids, we hypothesized that SSH2 facilitates Golgi-acrosome vesicular trafficking by modulating F-actin organization. To test this hypothesis, we performed immunostaining against PNA and GOPC (an acrosome-related protein implicated in vesicle trafficking from the Golgi apparatus to the acrosome (Yao et al., 2002)) in WT and *Ssh2* KO spermatids in seminiferous tubule squashes. In WT Golgi-phase spermatids, GOPC retained its distinct distribution, which was predominantly confined to the acrosomal region as indicated by its colocalization with PNA-positive structures near the nuclei (76.50 ± 3.70%). In contrast, in *Ssh2* KO spermatids GOPC exhibited dispersed localization in the cytoplasm and the extent of GOPC-PNA colocalization was significantly diminished (23.75 ± 7.72%) (Figure 6A, 6B). These results suggest that the participation of GOPC in proacrosomal vesicle trafficking is negatively affected in *Ssh2* KO mice.

**Fig. 6.**
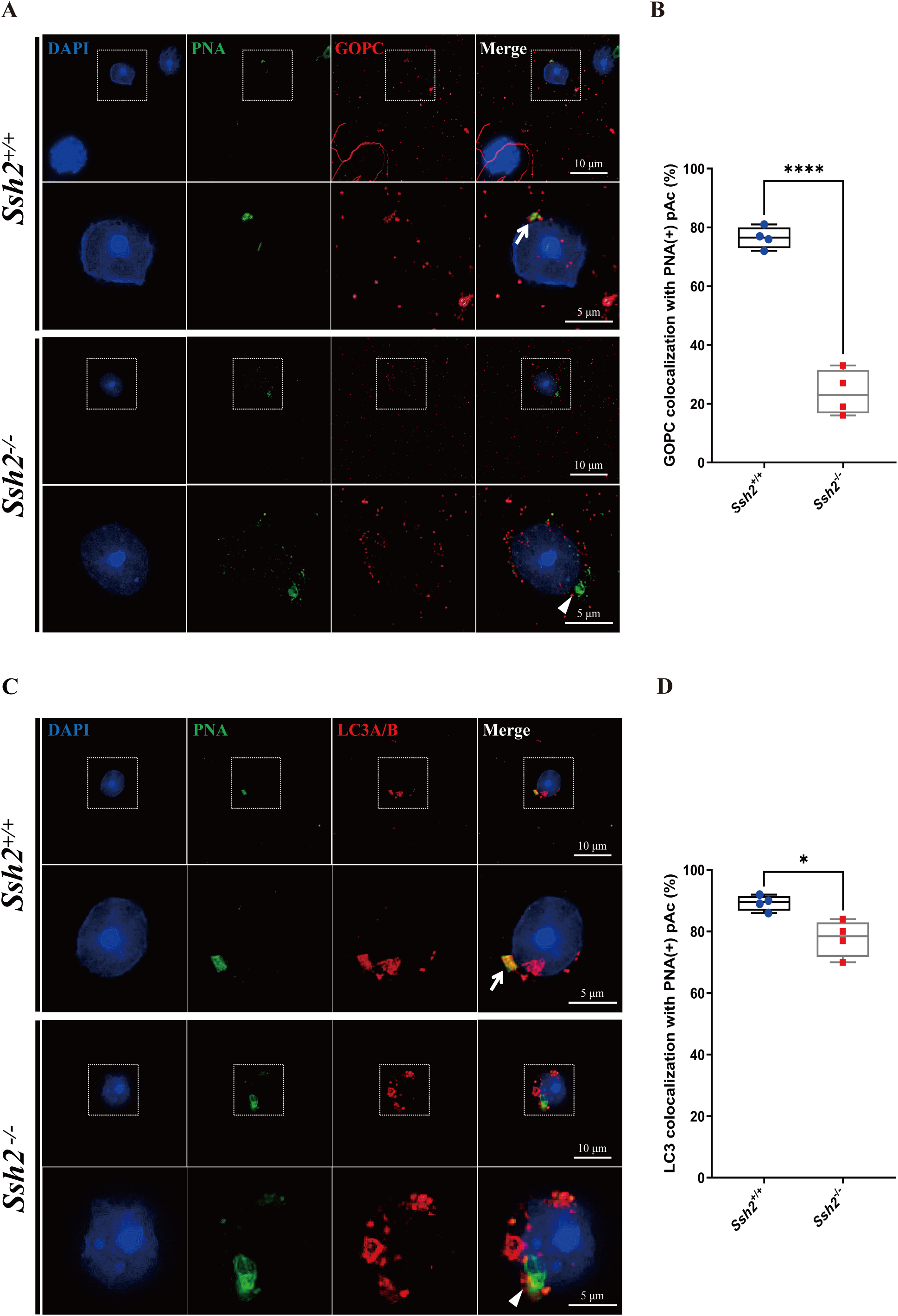
*Ssh2* KO spermatids exhibit defects in proacrosomal vesicle transport. **(A)** Immunofluorescence staining of the vesicular trafficking-related Golgi-specific protein GOPC (red) in WT and *Ssh2* KO round spermatids in testicular sections co-stained with PNA lectin (green). Nuclei were stained with DAPI (blue). Acrosomal debris was observed in *Ssh2* KO round spermatids. GOPC colocalized with PNA lectin in WT spermatids (upper panels, arrow indicates the co-localization) but not in those of *Ssh2* KO mice (lower panels, triangle indicates the absence of co-localization), as shown in the representative images. Framed areas are magnified beneath. Scale bars: original images, 10 μm; magnified images, 5 μm. **(B)** Quantitative analysis of GOPC and PNA lectin colocalization in WT mice, 76.50 ± 3.70%; *Ssh2* KO mice, 23.75 ± 7.72% (n = 4; 400 cells). Data are shown as mean ± SEM, \*\*\**p* < 0.001 by Student’s t-test. Bars indicate the range of data. **(C)** Immunofluorescence staining of the autophagosome-related protein LC3A/B (red) in WT and *Ssh2* KO round spermatids on the testicular sections co-stained with PNA lectin (green). Nuclei were stained with DAPI (blue). LC3A/B colocalized with PNA lectin in wild-type spermatids (upper panels, arrow indicates the co-localization) but not in those of *Ssh2* KO mice (lower panels, triangle indicates the absence of co-localization) as representative images showed. Framed areas are magnified beneath. Scale bars: original images, 10 μm; magnified images, 5 μm. **(D)** Quantitative analysis of LC3A/B and PNA lectin colocalization in WT mice, 89.45 ± 2.50%; *Ssh2* KO mice, 77.75 ± 5.91% (n = 4; 400 cells). Data are shown as the mean ± SEM, \*\*\**p* < 0.05 by Student’s t-test. Bars indicate the range of data. **Source data 6.** Observational datasets and original images.

We then tested whether the localization of the autophagy-related protein LC3, which is reported to function in Golgi-derived proacrosomal vesicle trafficking through its membrane conjugation (H. Wang et al., 2014), was affected by *Ssh2* KO.

Immunofluorescence analysis of LC3 was conducted on seminiferous tubule squashes from WT and *Ssh2* KO mice. We observed that LC3 was localized on proacrosomal vesicles in most of the WT spermatids (89.45 ± 2.50%), as indicated by its colocalization with PNA. In contrast, vesicle-like LC3-positive fluorescent signals surrounded the PNA-positive structures, and a portion of the LC3 molecules were not recruited to proacrosome vesicles in *Ssh2* KO spermatids (77.75 ± 5.91%) (Figure 6C, 6D). Together, these results indicate that *Ssh2* KO results in impaired trafficking of proacrosomal vesicles to the growing acrosome.

### *Ssh2* KO spermatids display impaired COFILIN phospho-regulation and thus have disturbed F-actin remodeling

In considering the potential molecular mechanisms underlying the observed impairment of acrosome biogenesis in *Ssh2* KO mice, we focused on the known role of SSH2 as a COFILIN phosphatase (Ohta et al., 2003). Given our observation of disorganized F-actin in *Ssh2* KO spermatids, we speculated that SSH2 acts as a modulator in actin remodeling though COFILIN dephosphorylation, which is essential for spermiogenesis. To pursue this further, we monitored the expression of COFILIN, phospho-COFILIN (p-COFILIN), and protein kinases that directly phosphorylate COFILIN (*e.g*., LIMK1 and LIMK2 (Arber et al., 1998)) by western blotting of testicular lysates of PD82 WT and *Ssh2* KO testes. The *Ssh2* KO mice had elevated levels of p-COFILIN compared to their WT littermates, whereas there were no significant differences in the levels of total COFILIN, LIMK1, or LIMK2 (Figure 7A, supplementary Figure S6). Accordingly, we also conducted an immunofluorescent analysis on testicular sections from adult WT and *Ssh2* KO mice using the antibody against p-COFILIN. Notwithstanding some mild nuclear staining in round spermatids, we did not observe evident p-COFILIN signals in the germ cells in WT testes. As expected, compared to WT mice, the signal intensity of p-COFILIN was dramatically increased in both the nucleus and cytoplasm of *Ssh2* KO spermatogenic cells, in particular in spermatids (Figure 7B). These findings suggest that COFILIN phospho-regulation is impaired in *Ssh2* KO testes with subsequent accumulation of p-COFILIN.

**Fig. 7.**
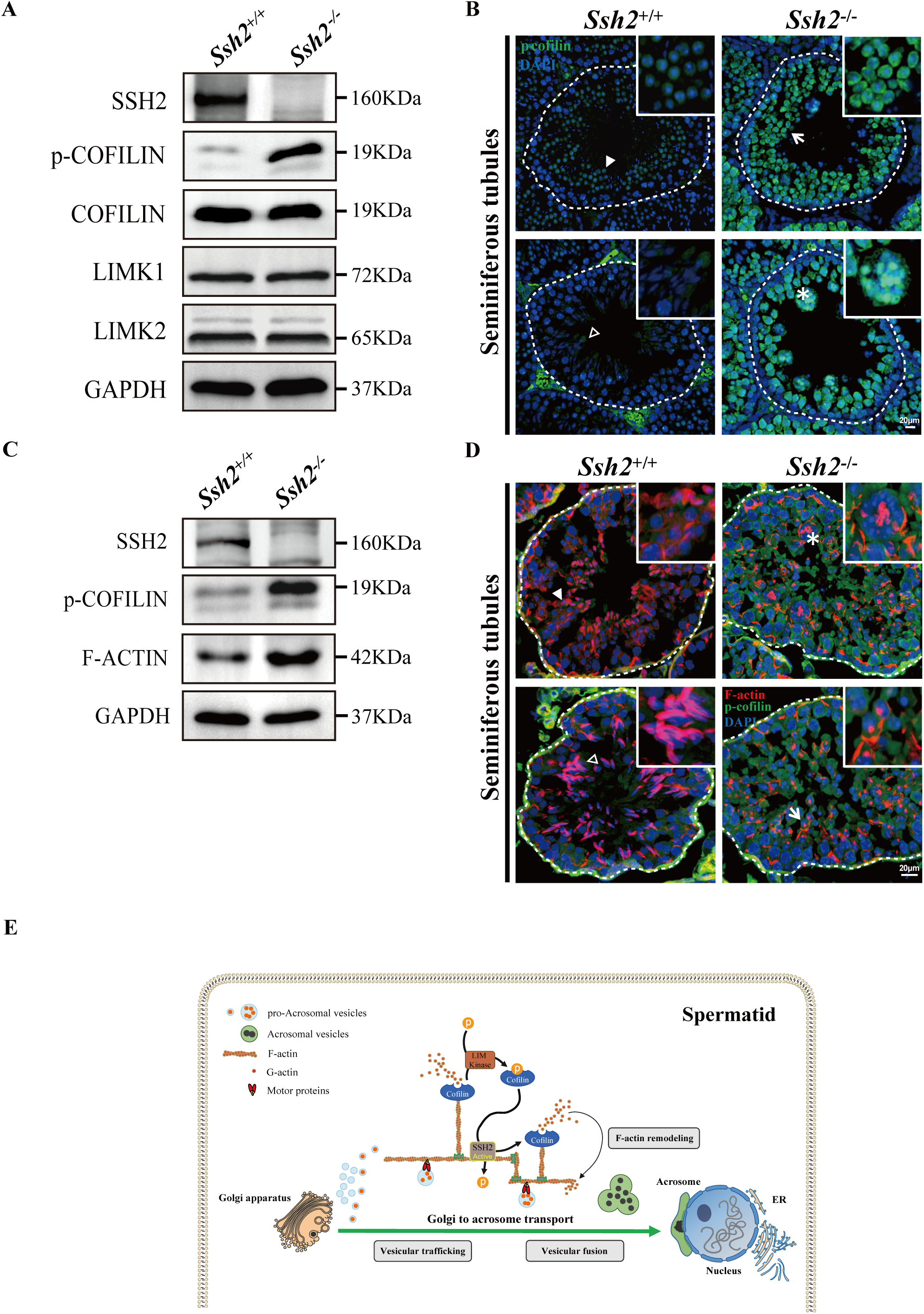
*Ssh2* KO spermatids display impaired COFILIN phospho-regulation that disrupts F-actin remodeling. **(A)** Immunoblotting against SSH2, p-COFILIN, COFILIN, LIMK1, and LIMK2 in WT and *Ssh2* KO testes sampled on PD82. GAPDH was used as the loading control. **(B)** Testicular sections of WT and *Ssh2* KO mice stained with p-COFILIN (green) and DAPI (blue), n = 3. The solid and hollow triangles point to the round spermatids and elongated spermatids of low p-COFILIN expression in seminiferous tubules of WT mice. The white arrow points to the round spermatids of strong p-COFILIN expression in seminiferous tubules of *Ssh2* KO mice. The asterisk indicates a germ cell cluster. Scale bars: 20 μm. **(C)** Western blot analysis of SSH2, p-COFILIN, and F-actin in lysates from 8-week-old WT and *Ssh2* KO testes, n = 3; GAPDH was used as the loading control. **(D)** Testicular sections of 8-week-old WT and *Ssh2* KO mice stained for p-COFILIN (green) and F-actin (red), n = 3. The solid and hollow triangles point to the round spermatids and elongated spermatids with intact F-actin organization in seminiferous tubules of WT mice. The white arrow points to the round spermatids of strong p-COFILIN expression with disorganized F-actin in seminiferous tubules of *Ssh2* KO mice. The asterisk indicates a germ cell cluster. Scale bars: 20 μm. **(E)** Proposed model for the functional role of the SSH2-COFILIN pathway in acrosome biogenesis. SSH2 participates in F-actin remodeling by regulating the phosphorylation of COFILIN. Intact and organized F-actin dynamics enables the transport of proacrosomal vesicles from the Golgi apparatus to the apical pole of the nucleus of the spermatid. F-actin also participates in the fusion of Golgi-derived vesicles and extra-Golgi vesicles. **Source data 7.** Original images/blots.

To further determine whether the effect of SSH2 on the actin cytoskeleton is mediated by the control of COFILIN phospho-regulation, we performed both western blotting and immunostaining against SSH2, p-COFILIN, and F-ACTIN in WT and *Ssh2* KO testicular lysates and sections, respectively. Not surprisingly, we observed increased expression of F-actin accompanied by p-COFILIN in *Ssh2* KO testes (Figure 7C). Fluorescence images also showed ubiquitous p-COFILIN signals overlapping with disorganized F-actin aggregates in *Ssh2* KO spermatids (Figure 7D). Collectively, these findings support a role for SSH2 in orchestrating F-actin remodeling through the control of COFILIN phosphorylation during acrosome biogenesis.

## Discussion

Although SSHs exert COFILIN phosphatase activity in mammals, the organ distribution of SSH family members varies significantly in mice, implying their similar but diverse biochemical functions (Ohta et al., 2003). In the present study, we show the essential role of SSH2 in acrosome biogenesis and male fertility based on the genetically engineered KO mice targeting *Ssh2*. Our findings demonstrate the reproductive phenotype of severe spermatogenic arrest at the early round spermatid stage with enhanced germ cell apoptosis, which leads to subsequent male infertility in *Ssh2* KO mice. The males lacking functional SSH2 display disrupted germ cell development starting from the spermiogenic step 2–3 round spermatids. Further examinations showed aberrant acrosome morphology in the mutant spermatids during spermiogenesis, characterized by fragmented acrosomal debris without the fusion of proacrosome vesicles. We also observed disorganized F-actin in the mutant developing spermatids accompanied by impaired proacrosomal vesicle trafficking, and this suggests a fundamental role for F-actin organization in acrosome formation. In addition, enhanced phosphorylation of COFILIN with thick F-actin fibers was seen in *Ssh2* KO testes. We interpret these findings as evidence for a role for SSH2 in acrosome biogenesis via the regulation of COFILIN-mediated F-actin remodeling (Figure 7E).

The implication of COFILIN in development has been described in model animals, as is evident by the embryonic lethality of *Cofilin-1* KO mice (Gurniak, Perlas, & Witke, 2005). Non-muscle COFILIN was identified as a component of tubulobulbar complexes in rat Sertoli cells (Guttman, Obinata, Shima, Griswold, & Vogl, 2004) and as a key activator of human sperm capacitation and the acrosome reaction (Megnagi, Finkelstein, Shabtay, & Breitbart, 2015), but its functions in mammalian spermatogenesis have remained largely elusive. Our finding that the enhanced cell apoptosis in *Ssh2* KO testes shares some similarities with the abnormal phenotype observed in the seminiferous tubules of *Limk2* KO mice (Takahashi et al., 2002) suggests a potential role for COFILIN in regulating the germ cell cycle in mice. Another line of evidence linking COFILIN to apoptosis comes from the investigations performed in cell contexts. Treating A549 cells with thapsigargin, an inhibitor of endoplasmic reticulum calcium ATPase, also induces apoptosis through the inhibition of COFILIN-mediated F-actin reorganization (F. Wang et al., 2014). Elevated COFILIN phosphorylation after SSH2 knockdown was previously demonstrated to induce the activation of Caspase3/7 in a human renal cell carcinoma cell line (Lu et al., 2014), and activation of Caspase3/7 is known to trigger apoptotic cell death (Saller et al., 2010). Consistently, we observed distinct Caspase-3 activation accompanied by correlative changes in the expression of Bcl-2 family members in *Ssh2* KO testes. Based on these findings from published literatures and our current study, we speculate that the mutant spermatids in *Ssh2* KO mice possibly undergo cell cycle arrest via the Bcl-2/Caspase cascade due to impaired COFILIN phosphorylation. Future studies are needed to further elucidate the mechanisms underlying this actin-based process.

Owing to the resolution limitation of microscopy, we could not directly image the ultrastructure of F-actin tracts between the Golgi body and the acrosome. However, the observation from TEM that the lack of the electron-dense acroplaxome, a cytoskeletal actin-rich scaffold plate that anchors the acrosome to the nuclear envelope (Kierszenbaum, Rivkin, & Tres, 2003a), in *Ssh2* KO spermatids indicates a potential role of SSH2 in the shaping of the acroplaxome. Notably, defective acroplaxome structures with impaired acrosome biogenesis were observed in mutant spermatids lacking homeodomain-interacting protein kinase 4 (Crapster et al., 2020), which is responsible for the regulation of F-actin remodeling during spermiogenesis. We infer from these findings that acrosome biogenesis in *Ssh2* KO mice might also be affected by the acroplaxome defects that result from disordered F-actin remodeling, and further experiments should be performed to confirm this.

In conclusion, our study confirms the requirement for SSH2 during mammalian spermatogenesis, specifically in acrosome biogenesis. Because the Slingshot family are evolutionary conserved in mammals, we believe that some mutations in the *SSH2* gene should exist in non-obstructive azoospermia, although no such mutations have, to the best of our knowledge, been reported. At the very least, clues to elucidating the phenotypes of the *Ssh2* KO mouse model can help us to understand the pathogenic mechanisms of this potential factor in causing azoospermia.

## Author Contributions

H.L., J.M., and Z-J.C. performed study concept and design; K.X., X.S., and Y.L. performed the development of methodology and investigation; X.S., Y.L., H.L., K.F., and R.W. analyzed the data; H.L., G.L., W.C., and J.M. supervised the article; K.X., and X.S. wrote the manuscript; X.S., and H.L. reviewed and revised the manuscript.

## Data Availability

All data generated or analysed during this study are included in the manuscript and the Source data files as:

Figure 1-source data 1

Figure 2-source data 2

Figure 3-source data 3

Figure 4-source data 4

Figure 5-source data 5

Figure 6-source data 6

Figure 7-source data 7

Figure supplement S1-source data 8

Figure supplement S2-source data 9

Figure supplement S3-source data 10

Figure supplement S4-source data 11

Figure supplement S5-source data 12

Figure supplement S6-source data 13

## Ethics approval

All procedures were in accordance with the ethical standards approved by the Animal Use Committee of the School of Medicine, Shandong University, for the care and use of laboratory animals. The research was approved by the Institutional Review Board of Shandong University.

## Acknowledgments

We thank our colleagues at the Center for Reproductive Medicine, Shandong University, for their technical support.

## Supplementary Information for

**Fig. S1.**
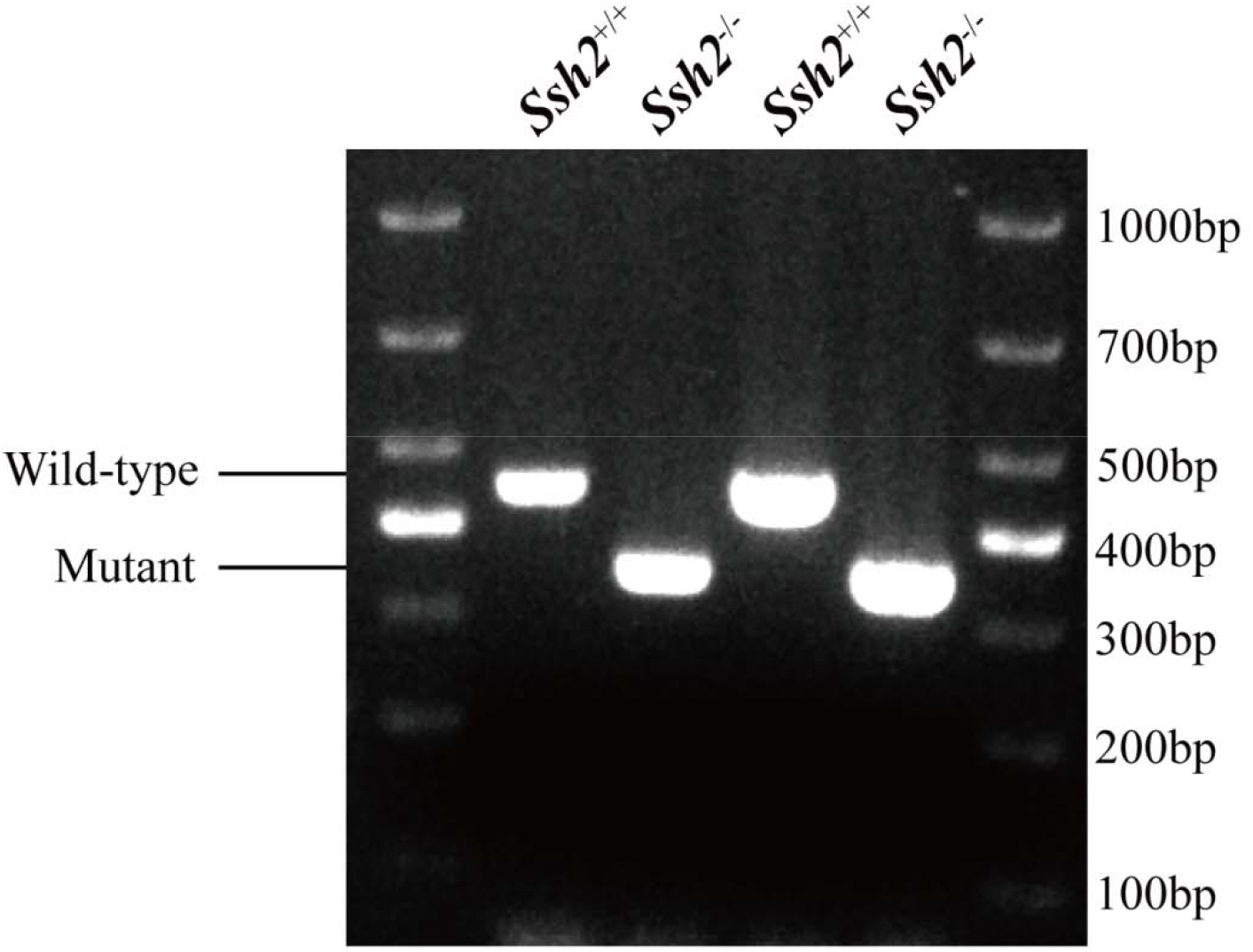
PCR genotyping for WT and *Ssh2* KO mice. PCR genotyping demonstrating the absence of the WT band in *Ssh2* KO mice. **Source data 8.** Original gels.

**Fig. S2.**
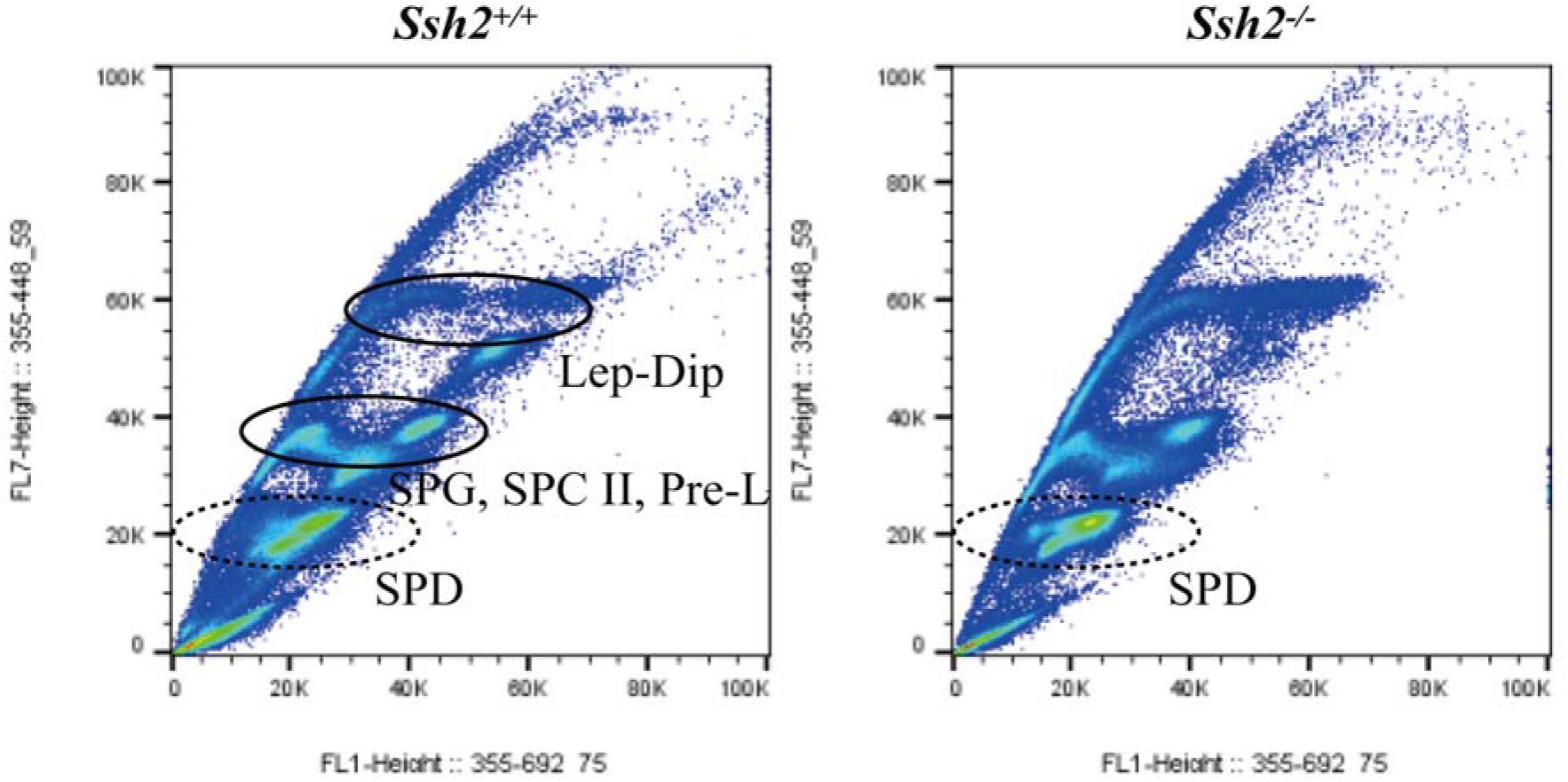
FACS assessment in spermatogenic cells from WT and *Ssh2* KO murine testes. Distribution of spermatogenic cells by FACS. The number of spermatids (round/elongated spermatids in WT testes and round spermatids in *Ssh2* KO testes) showed no significant variation in *Ssh2* KO testes (as the dotted circles indicate). Lep, leptotene spermatocyte; Dip, diplotene spermatocyte; SPG, spermatogonia; SPC II, secondary spermatocyte, Pre-L, pre-leptotene spermatocyte; SPD, spermatid. **Source data 9.** Original images.

**Fig. S3.**
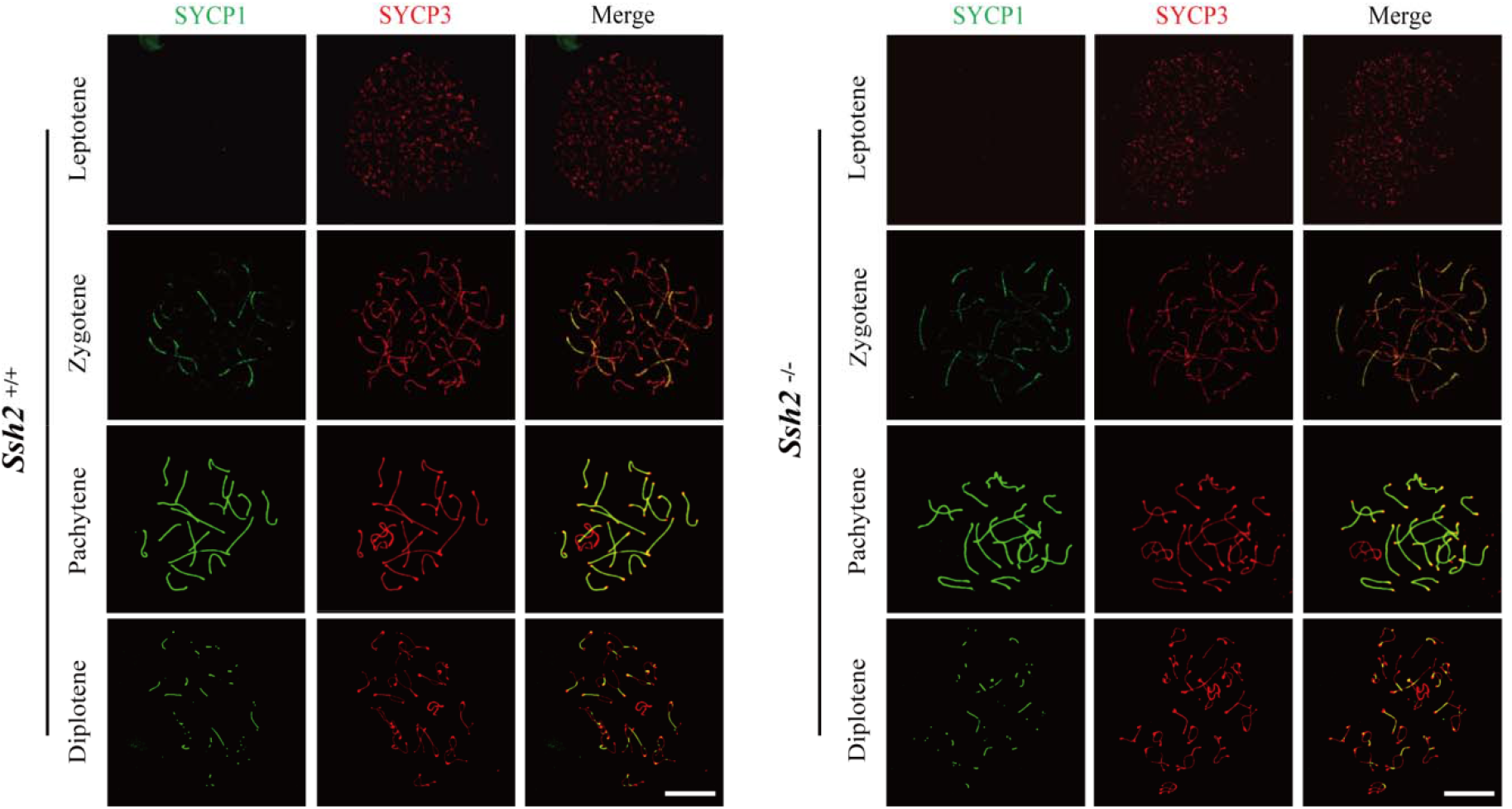
S3 SSH2 is not required for meiotic progression in mouse spermatogenesis. Testes from nearly 3-week-old WT mice (left panels) and *Ssh2* KO mice (right panels) were sampled for preparing chromosome spreads immunostained for SYCP1 and SYCP3. Spermatocytes from *Ssh2* KO mice at leptotene, zygotene, pachytene, and diplotene of prophase I show no obvious abnormalities. Images are representative of three mice per genotype. Scale bars: 10 μm. **Source data 10.** Original images.

**Fig. S4.**
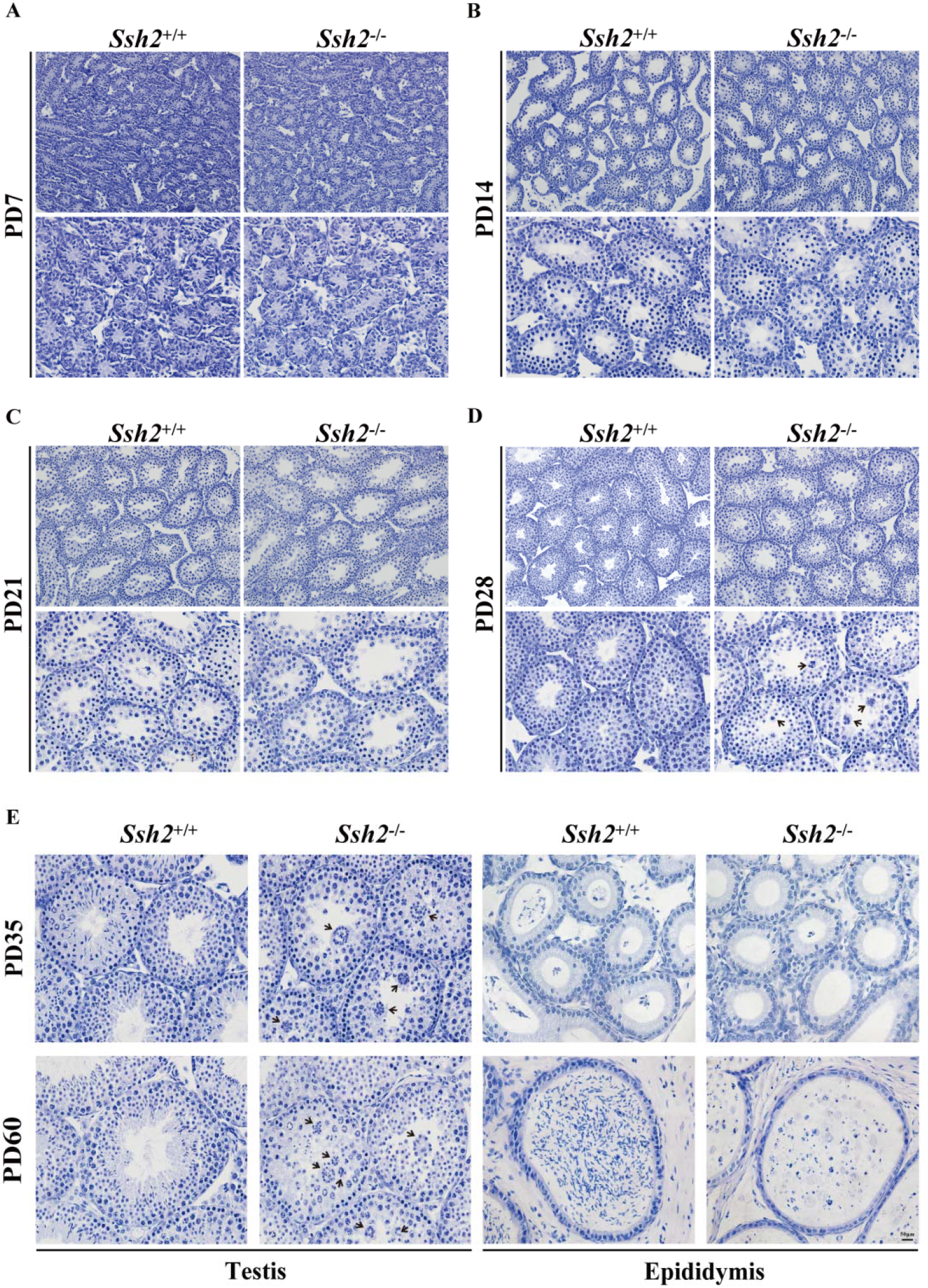
Comparison of spermatogenesis in WT and *Ssh2* KO mice. Comparison of spermatogenesis in WT and *Ssh2* KO mice assessed in testicular sections stained for hematoxylin at PD7, PD14, PD21, PD28, PD35, and PD60 accompanied by epididymal sections at PD35 and PD60. Black arrows: round spermatid clusters. Scale bar: 50 μm. **Source data 11.** Original images.

**Fig. S5.**
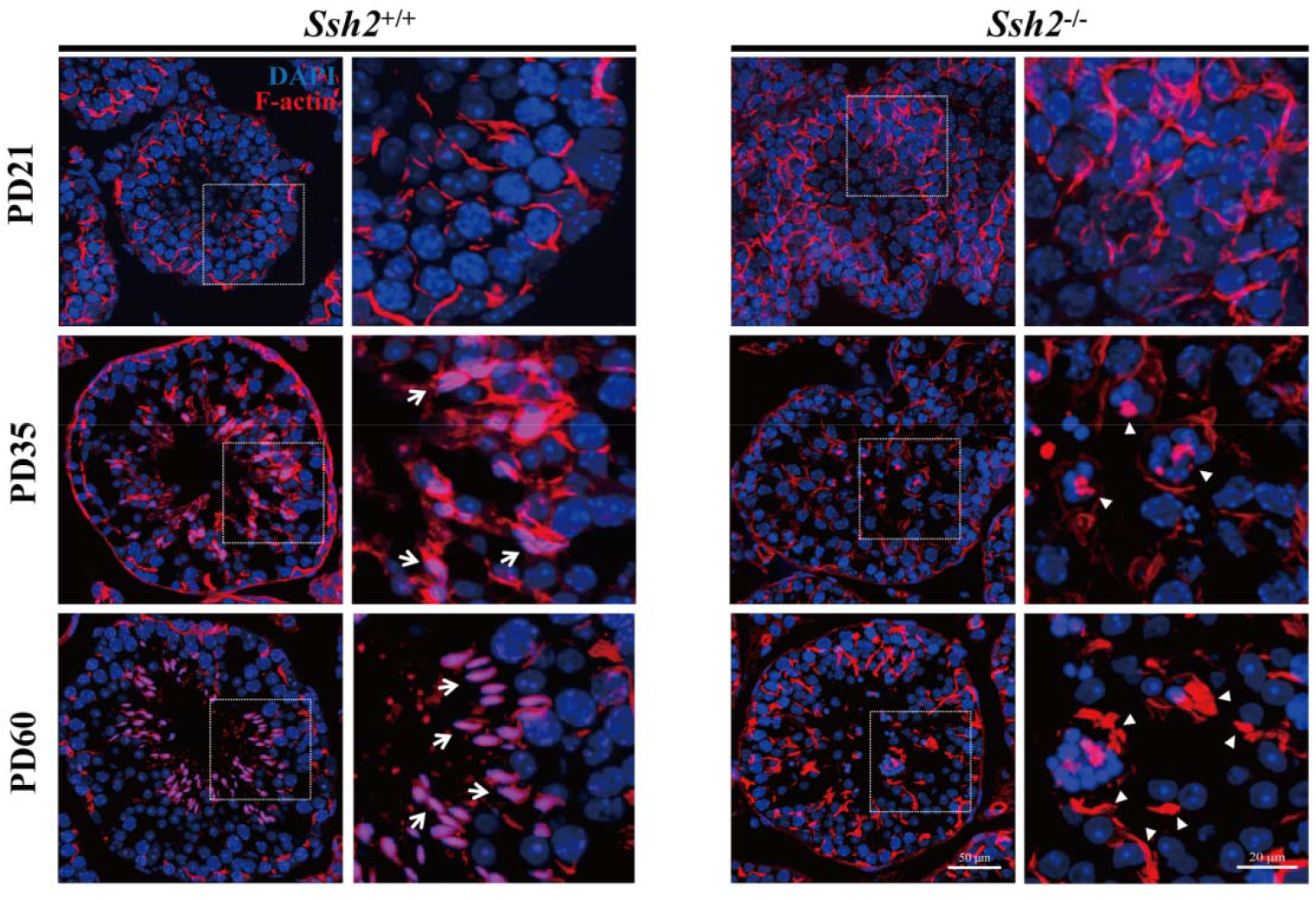
Disordered F-actin in *Ssh2* KO testis. Fluorescence imaging of phalloidin-labeled F-actin (red)-stained seminiferous tubules from WT (left panels) and *Ssh2* KO (right panels) murine testes at PD21, PD35, and PD60. Nuclei were stained with DAPI (blue). Representative images show the sharp sickle head of WT spermatids “hooped” by F-actin bundles (indicated with arrows), whereas actin filaments aggregated to form several lumps in *Ssh2* KO spermatids (triangles). The framed regions are shown at higher magnification to the right. Scale bars: Original images, 50 μm; magnified images, 20 μm. **Source data 12.** Original images.

**Fig. S6.**
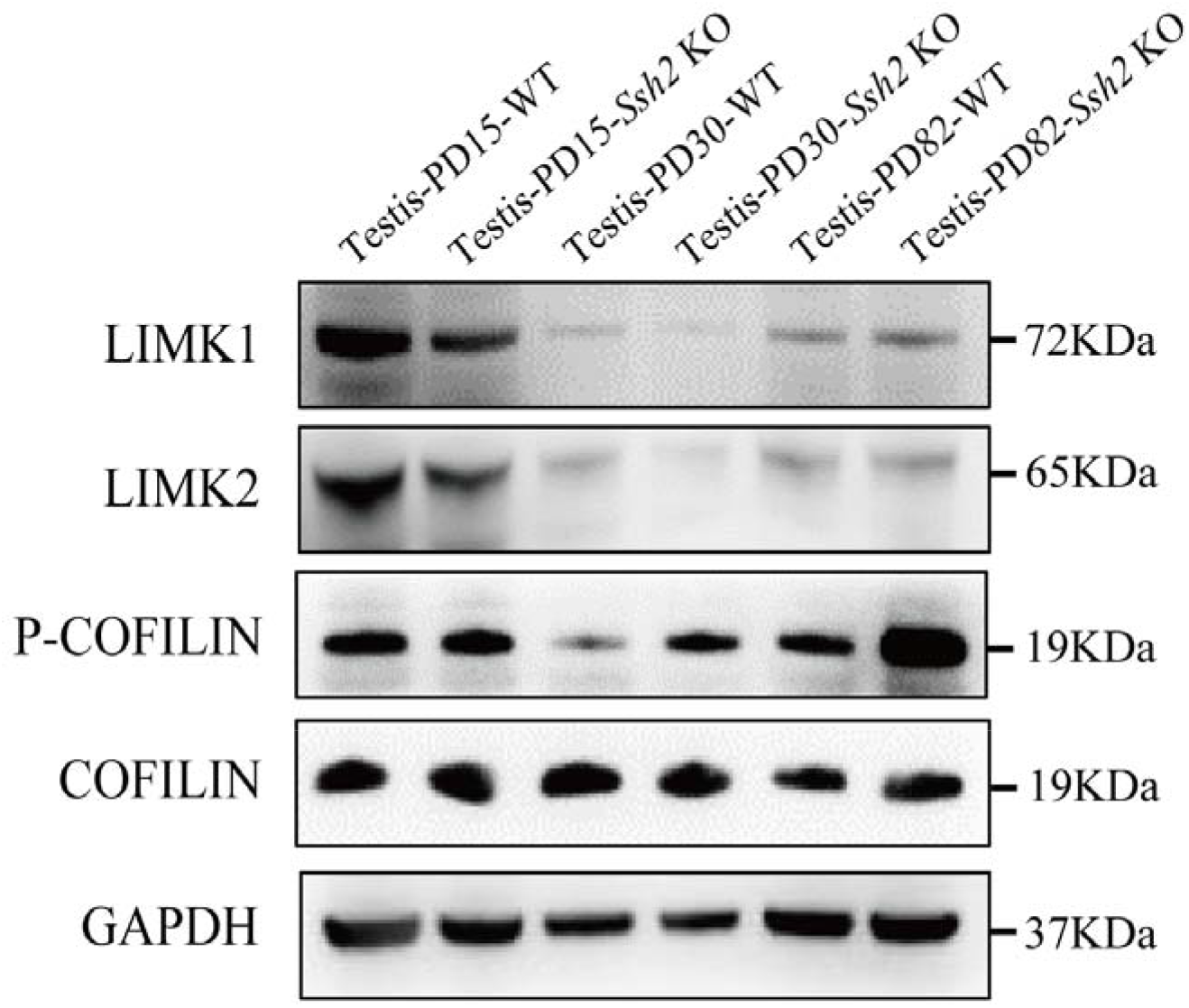
Cofilin-associated protein expression level in WT and *Ssh2* KO testes. Protein extracts from testes isolated from WT and *Ssh2* KO mice at PD15, PD30, and PD82 were used for western blot analysis for LIMK1, LIMK2, p-COFILIN, and COFILIN. GAPDH was used as the loading control. **Source data 13.** Original blots.

